# Compensatory hippocampal neurogenesis in the absence of cognitive impairment following experimental hippocampectomy in adult rats

**DOI:** 10.1101/2020.06.04.135350

**Authors:** Giuliana T. M. Cardoso, Walace Gomes-Leal, Edna C. S. Franco, Jessica S. Gama, Francinaldo L. Gomes, Ana Leda F. Brino, Antonio Pereira, Silene M. A. Lima.

## Abstract

Temporal lobe epilepsy (TLE) is the commonest type of focal epilepsy in adult humans. In refractory TLE, patients are indicated for unilateral resection of the affected hippocampus (hippocampectomy), which generally does not cause any cognitive impairment. Once adult hippocampus is a region of endogenous neurogenesis, we have hypothesized that a compensatory increase in hippocampal neurogenesis might occur in the remaining hippocampus after unilateral hippocampectomy. To test this hypothesis, we performed unilateral hippocampectomy in adult Wistar rats (n=12). Sham animals were not hippocampectomized (n=6). Animals were deeply anesthetized and adjacent cortex and hippocampus of the left hemisphere were completely removed. They were perfused at 15 (G15, n=6) or 30 (G30, n=6) days post-surgery. Behavioral tests were performed to address possible cognitive impairments. We did not find any cognitive impairment in the hippocampectomized animals. Histopathology was performed using thionine staining and mature neurons and migratory neuroblasts were immunolabeled using anti-NeuN and anti-doublecortin (DCX) antibodies, respectively. The remaining hippocampus presented higher numbers of DCX positive cells compared to control (p<0.001) at both G15 and G30. The results suggest increased compensatory adult neurogenesis following experimental unilateral hippocampectomy in adult rats, which may contribute to absence of cognitive impairments.

## 1. Introduction

Epilepsy is a chronic neurological disorder affecting nearly 1% to 2% of the World population. Its hallmark is an abnormal increase in the predisposition to seizures (Fisher et al., 2005). Among epilepsies, temporal lobe epilepsy (TLE) is the most common type of refractory epilepsy in adult humans. In TLE, epileptic focus is located in the temporal lobe, and, in about 50% of cases, in the hippocampus (Anyanwu et al., 2018). Unilateral amigdalohippocampectomy is often effective in seizure control.

It has been widely reported that bilateral damage to hippocampal formation may produce impairments in the capacity of memory consolidation in humans, and that the extent of such damage is related to the extent of damage to hippocampal formation and adjacent cortical areas (Squire, 2009; Squire & Wixted, 2011). However, patients with TLE submitted to unilateral hippocampectomy usually do not present cognitive deficits (Anyanwu & Motamedi, 2018; Jobst & Cascino, 2015; Vojtěch et al., 2012). There is a continuous production of new neurons in the adult hippocampus both in experimental animals (Kempermann, 2015; Lima and Gomes-Leal, 2019) and in the human brain, even in elderly people (Boldrini et al., 2018; Lima and Gomes-Leal, 2019; Moreno-Jiménez et al., 2019). It follows, that the existence of a compensatory neurogenic mechanism might underlie the reason by which postoperative patients generally do not present cognitive deficits related to memory consolidation.

To test this hypothesis, we performed experimental hippocampectomy in adult rats to investigate possible compensatory neurogenic events in contralateral side to surgery, as well as whether the procedure causes some kind of learning and memory deficits in the surged subjects.

## 2. Material and Methods

### 2.1 Animals

18 adult male adult Wistar (*Rattus norvegicus*) rats were used. Procedures were evaluated and approved by the Ethic Committee on Research with Experimental Animals of the Federal University of Pará (CEUA-UFPA) (approval number 6552280616). Procedures for the use of experimental animals suggested by Society for Neuroscience, US National Research Council’s Guide for the Care and Use of Laboratory Animals, US Public Health Service’s Policy on Humane Care and Use of Laboratory Animals, Guide for the Care and Use of Laboratory Animals and CEUA-UFPA were followed.

Animals were maintained in the animal facility at the laboratory of Neuroprotection and Experimental Neuroregeneration in individual cages with controlled temperature at 23°C, light/dark cycle of 12 hours and water and feed *ad libitum*.

To test the raised hypothesis, three experimental groups were delineated: animals submitted to surgical removal of the left hippocampus (unilateral hippocampectomy) with a survival time of 15 (G15) and 30 (G30) days. This period was chosen based on the time of neuroblast formation and neuronal maturation in rodents). Animals in the control group (Gc) were not submitted to any surgical procedures (sham animals).

### 2.2 Surgical procedures

Animals were anesthetized with intraperitoneal injections of xylazine hydrochloride (10mg/kg) and ketamine hydrochloride (80mg/kg). Time required for abolition of both corneal and interdigital reflex was waited for the surgical procedure to begin. When necessary, anesthesia reinforcement injections were administered with a dose adjusted to 1/3 of the initial dose.

The animal was fixed in the stereotaxic apparatus and asepsis performed. A longitudinal surgical incision was made for exposure of the skull and visualization of bregma and sagittal sutures. Coordinates for the location of the dorsal hippocampus were adjusted according to the stereotaxic atlas by Paxinos and Watson (1998). We used the following stereotaxic coordinates in relation to the bregma: −1.5 mm in the mid-lateral direction (M/L) and +2.8 mm in the anterior-posterior direction (A/P). With the use of a dental drill, craniotomy was performed in the region around the hippocampus reference area (indicated by the stereotaxic coordinates). With the aid of a stereo surgical microscope, meninges and adjacent cortex were removed for exposure of the dorsal hippocampus, which was individualized and removed in block. Hemostatic powder was used to minimize bleeding. At the end of the procedure, skin and muscles were sutured and an anti-inflammatory ointment was administered to assist the healing process. Antibiotic was administered to the animal for three days at 24-hour intervals.

### 2.3 Behavioral procedures

An 8-arm radial maze was used to evaluate whether the surgical intervention produces learning and memory deficits in the experimental animals. The eight arm Radial Maze consists of an octagonal platform with a diameter of 34 cm from which eight arms, 47.5 cm long and 13 cm wide, are placed at a height of 60 cm from the ground. Before the start of the test, animals were submitted to a food adaptation period (around 4 days) in which they had a decrease in the amount of feed. At 24 hours before the test onset, animasl were submitted to food deprivation.

The behavioral procedures were initiated in the experimental groups G15 and G30 on the eighth (8th) and twenty-third (23rd) days, respectively, after the hippocampal removal surgery so that the behavioral test was ended at the tenth fifth and thirtieth day after surgery. Animals in Gc group were not submitted to surgical procedures.

Animals were exposed for 7 days to the maze. During this period, rats explored the maze (once a day per animal). Every day, four specific arms of the maze were baited with a piece of rat food, always in the same four arms. For each exploration day, animal was placed on the central platform and allowed to freely explore the apparatus. The exploration time was set as ten minutes (600 seconds) or the time required for the animal to eat all the food present in the apparatus.

To assess learning during exploration, total duration of each session was quantified over 7 seven days of exposure to the maze. The reference memory measure was evaluated by the number of entries into the arms without food. Each of these entries was counted as a reference memory error (RME). The measure of working memory was the amount of entries into the arms in which the rats had already consumed the food. Each of these re-entries was counted as a working memory error (WME). These two measures (RME and WME) were registered only in the last two days of exposure to the maze.

### 2.4 Perfusion and preparation of biological material

After survival times of 15 and 30 days, animals were deeply (*i.p*) anesthetized with xylazine hydrochloride (10mg/kg) and ketamine hydrochloride (80mg/kg). After reflex abolishment, animals were transcardially perfused with heparinized 0.9% phosphate-buffered saline (PBS) followed by 4% paraformaldehyde solution in 0.1M PB.

Brains were removed, immediately immersed in 4% paraformaldehyde solution for 72 h and cryoprotected for 6 days in different gradients of sucrose-glycerol solutions in 0.1M phosphate Buffer. Brains were then frozen in Tissue Tek and coronal sections were cut at 30μm thickness using a cryostat (Leica CM1850).

### 2.5 Nissl staining

To evaluate gross histology, representative sections from, Gs, G15 and G30 animal groups were mounted onto gelatinized slides and submitted to Nissl staining. In short, slides were immersed in decreasing concentration of alcohol solutions (100%, 70%, 50% and 20% alcohols) in distilled water for 1 minute each, followed by immersion in 0.1% thionin solution for 8 seconds. Slides were then immersed in distilled water and in increasing concentration of alcohol solutions (alcohol 20%, 50%, 70%, 90% and 100%) with incubation time of 45 seconds each. They were then immersed in xylol for 5 minutes, mounted in Permount® and coversliped for further analysis.

### 2.6 Immunolabeling protocols

To immunolabel cell bodies of differentiated neurons, we used an antibody against NeuN, a protein present in the nucleus of mature neurons (Mullen et al, 1992). To immunolabel immature neurons (neuroblasts), we used an antibody against, doublecortin, a microtube-associated protein used as a specific marker for neuroblasts (Gleeson et al., 1999).

#### 2.6.1 Anti-NeuN immunolabeling protocol

Details of the immunolabeling protocol were reported in our previous publications (Thored et al., 2009). In short, sections were transferred to 24 wells culture plates, rinsed three times in 0.1M PBS (5 min each) and incubated with 3% hydrogen peroxide solution for 15 min. Another wash battery in 0.1M PBS 0.1M was performed. Sections were incubated in citric acid solution in a water bath at 90 °C for 20 min and further allowed to cool for about 10 min at room temperature (20 °C). Another wash battery with 0.1MPBS was performed. Tissue permeabilization was performed with saponin solution for 10 min, followed by another washing battery with 0.1M PBS. Tissue was then incubated with 10% BSA solution for 30 min and subsequently incubated with primary antibodies overnight (anti-NeuN, LSBio® LS-C312122, 1:50).

On the second day, sections were labeled by LSAB2 system-HRP (Dako® - K0675). They were rinsed three times with 0.1M PBS to remove excess of primary antibody (5 min each), and subsequently incubated with biotin for 30 min at 37 °C, followed by another wash battery with PBS 0.1M before the incubation with streptavidin for 30 min at 37 °C. Another washing battery with 0.1M PBS was made to remove streptavidin excess, and then, the revelation was made using the liquid DAB + substrate chromogen system (Dako® - K3468). The reaction time ranged from 04 to 05 min and the reaction was stopped by washing in 0.1M phosphate buffer solution.

#### 2.6.2 Anti-DCX immunolabeling protocol

On the first day, sections were transferred to 24 well culture plates as for the previous antibody. They were rinsed twice in 0.1M PBS (5 min each rinse), pretreated in 0.2M boric acid (pH 9.0) at 60 °C for 25 min and allowed to cool for 20 min at room temperature (20 °C). Sections were then incubated with 3% hydrogen peroxide in methanol solution for 20 min, rinsed 3 times with 0.05% PBS/TWEEN (5 min each) and incubated with 10% normal horse serum in PBS for one hour. They were then incubated with primary antibody overnight (Anti-DCX, Santa Cruz Biotechnology Cat# sc-8066, RRID:AB_2088494, 1:100). On the second day, sections were rinsed 3 times in PBS/TWEEN, and incubated with secondary antibody biotinylated horse-anti-goat (Vector Laboratories®, 1:100) for 2 h at room temperature. Another PBS/TWEEN washing battery was performed and sections were then incubated in an avidin-biotin-peroxidase complex (ABC kit, Vector Laboratories®) for 2 h. Sections were then rinsed 4 times in PBS /TWEEN (5 min each) and DAB reacted with reaction time ranging from 4 to 5 min. The reaction was stopped by washing sections in 0.1M phosphate buffer solution.

All sections were mounted onto gelatinized slides, dehydrated in alcohol gradients, cleared in xylol, mounted onto slides using Permount and coversliped for further analysis.

### 2.7 Qualitative analysis

Sections of all the experimental groups were surveyed using a Zeiss Imager Z1 optical microscope. Representative images were obtained using a digital camera (Zeiss AxioCam HRc) attached to the microscope.

### 2.8 Quantitative analysis

Quantitative methods were based on our previous reports for hippocampal countings (Silva et al., 2013). DCX positive cells from all experimental groups were counted using a graticule attached to the eyepiece of an optical microscope (Nikon, 50i). A 40X objective was used. The graticule counting area was 0.0625mm^2^. At least 4 sections were counted per animal.

Countings were performed along the full extent of the dentate gyrus and the initial area of the hippocampal hilus. In the dentate gyrus, graticule was positioned in the granular cell layer. Due to compact morphology of this hippocampal layer, counting area was set as 0.01875mm^2^, a fraction of the whole graticule area.

### 2.9 Statistical analysis

Kruskal Wallis test with Dunn post-test or one-way-ANOVA with Tukey post-test were used for comparisons between groups. The significance level was set at p<0.01 and the analysis were performed using the GraphPad Prism 7.0 software.

## 3. Results

### 3.1 Left hippocampus was completely removed after surgical procedure

Nissl staining revealed the normal hippocampal morphology in both control (Figure 1A) and surged animals (Figure 1B). Hippocampectomy was successfully achieved, as the left hippocampus was completely removed after surgery while the right hippocampus was preserved (Figure 1B).

**Figure 1.**
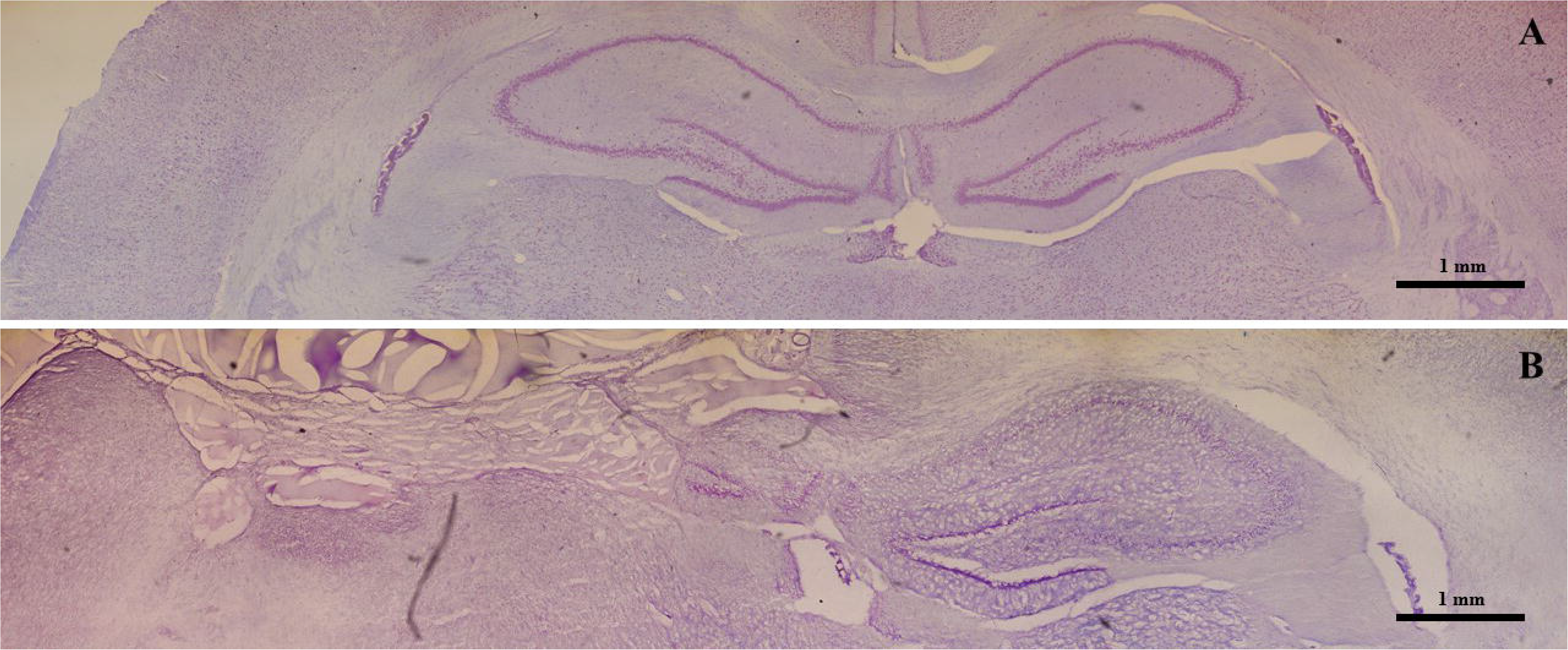
Effective experimental unilateral removal of rat hippocampus. Photoreconstruction of a Wistar rat brain stained with 0.1% thionin. A: Coronal section obtained from a control animal presenting the right and left intact hippocampus. B: Section obtained from an animal submitted to a left hippocampectomy. Note the absence of the left hippocampus.

### 3.2 Increased numbers of migratory neuroblasts in the hippocampus of animals submitted to unilateral hippocampectomy

The immunolabeling of migratory neuroblasts was performed using an anti-DCX antibody, which revealed DCX positive cells (DCX+) in the dentate gyrus and hilus of animals belong to all experimental groups. Nevertheless, quantitative differences in the number of DCX+ cells were observed between groups (Figure 2). There was an increase in the number of DCX+ cells in groups G15 (Figure 2B and 2E) and G30 (Figure 2C and 2F) compared to Gc (Figure 2A and 2D). These results were quantitatively confirmed (Figure 2G-H).

**Figure 2.**
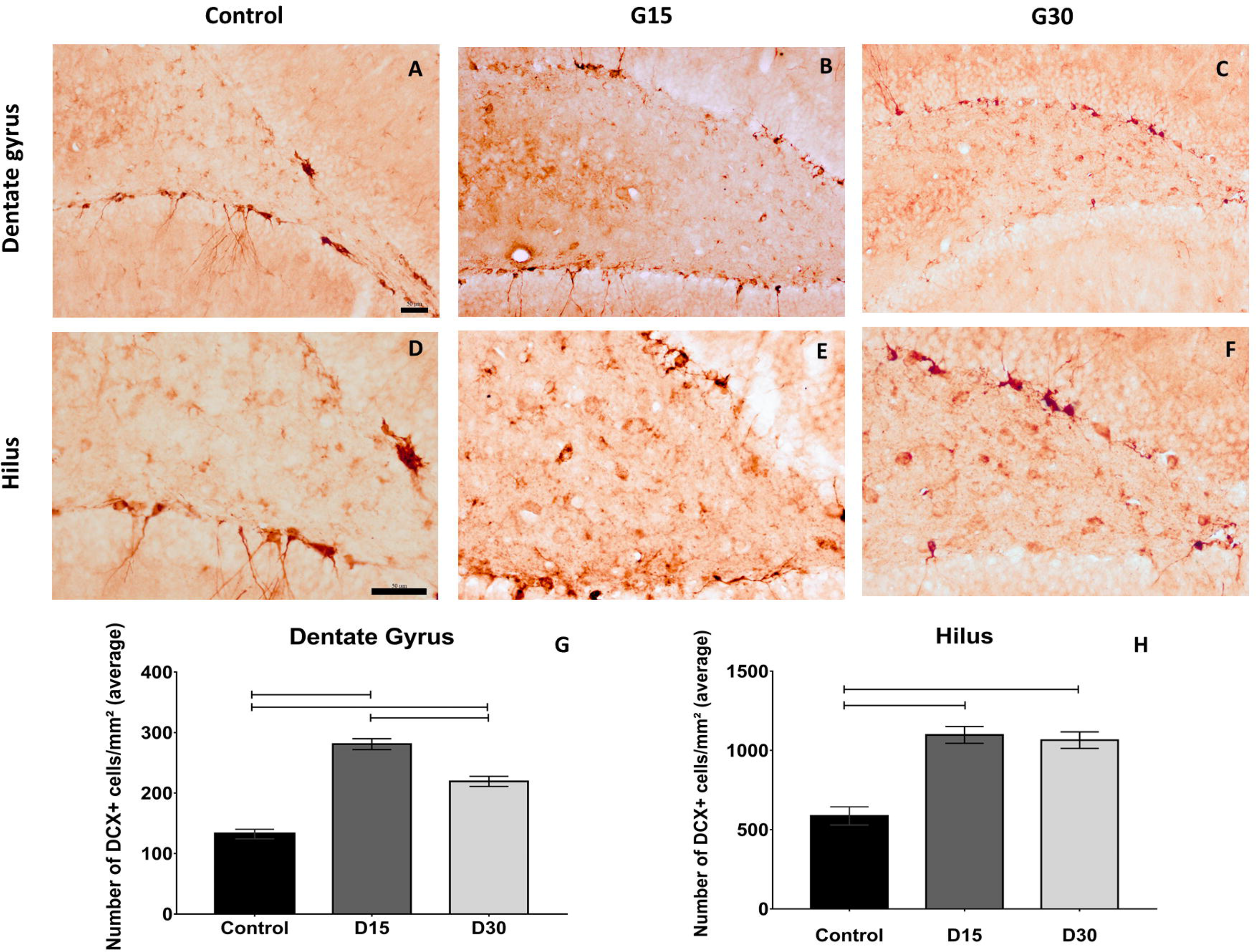
Compensatory increase in rat hippocampal neurogenesis after unilateral hippocampectomy. Immunolabeling for DCX in coronal sections from animals belonging to all experimental groups (Control, G15 and G30). There was increase in the number of neuroblasts in both dentate gyrus (B-C) and hilus (D-F) compared to control (A, D). This has been confirmed by quantitative analysis (G, H, Kruskal Wallis with dunn posthoc test (dentate gyrus); one-way-ANOVA with Tukey post-test-test (hilus), p<0.01). Scale bars: 50μm; A-C (objective 20X); D-F (objective 40X.).

Statistical comparisons between groups revealed that numbers of DCX+ cells were higher in G15 and G30 compared to Gc (Kruskal walis/with Dunn’s posthoc test, p <0.0001, Figure 2G-2H). Comparisons between surgical groups revealed that numbers of DCX+ cells in the dentate gyrus were higher in G15 compared to G30 (Figure 2G, p <0.0001,). This was not the case in the hilus (p >0.0001, Figure 2H). In this region, numbers of DCX + cells remained equally elevated in both survival times. These results indicate increased numbers of neuroblasts in the remaining hippocampus after experimental unilateral hippocampectomy.

### 3.3 Hippocampal cytoarchitecture and morphology after hippocampectomy

The immunolabeling of mature neurons was performed using an anti-NeuN antibody, which revealed the conspicuous hippocampal cytoarchitecture and morphology in all experimental groups (Figure 3). A neuroanatomical analysis was performed in order to infer some alteration on the normal hippocampal cytoarchitecture and morphology in the contralateral hippocampus after surgery.

**Figure 3.**
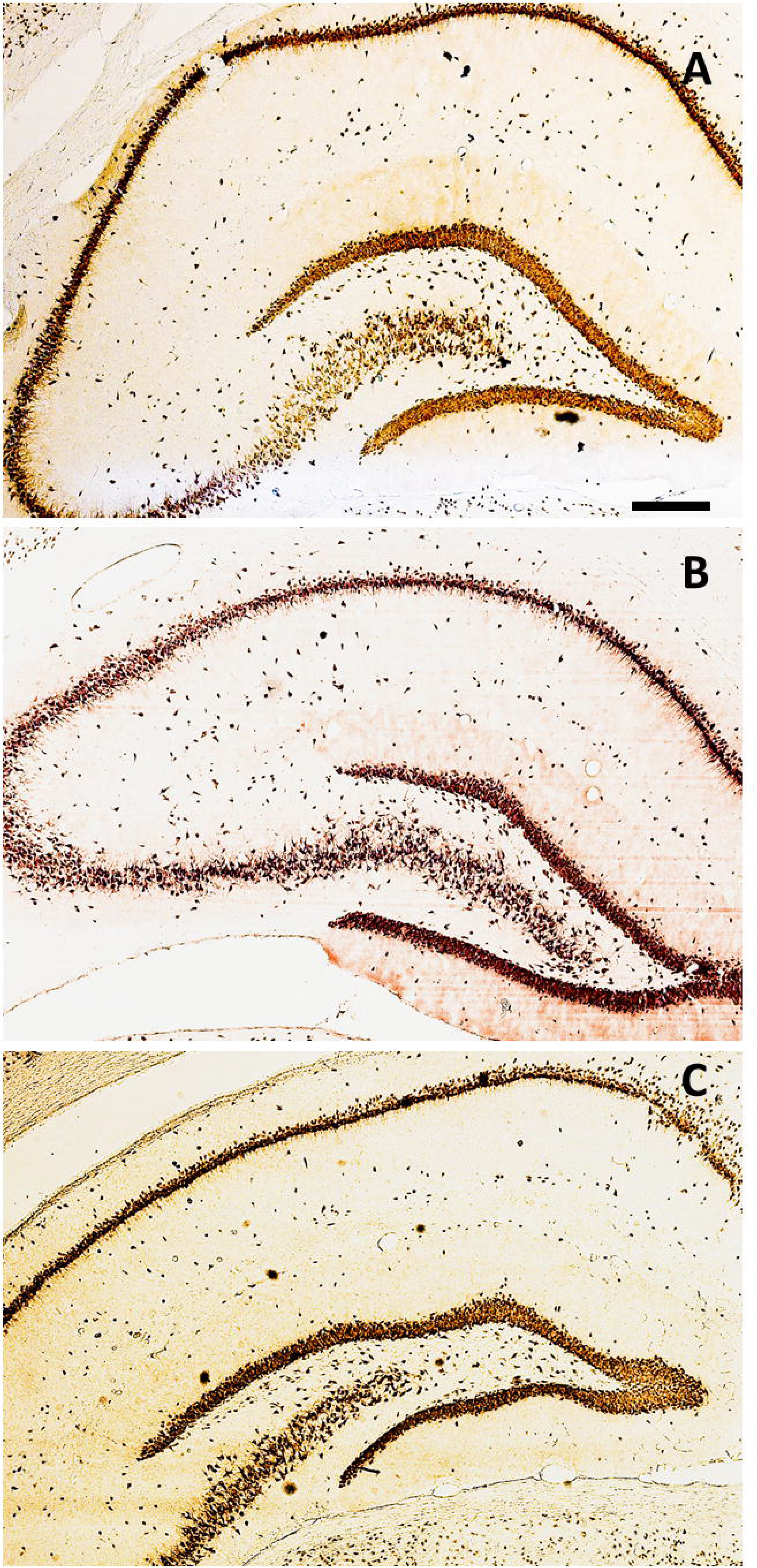
Distribution of NeuN+ cells along hippocampal areas. Coronal sections immunolabelled for NeuN in control group (A), G15 (B) and G30 (C). Scale bar = 300μm.

In CA1, NeuN immunolabeling revealed cell bodies of mature neurons with normal cytoarchitecture for all experimental groups (Figures 4A-F). In CA3 (Figure 4G-L), NeuN immunolabeling showed a differential distribution of mature neurons in the contralateral hippocampus after surgery. It was noted a thicker layer and higher number and of NeuN + cells in G15 (Figures 4H, K) compared to control (Figures 4G,J), which was not evident in G30 (Figures 4I, L).

**Figure 4.**
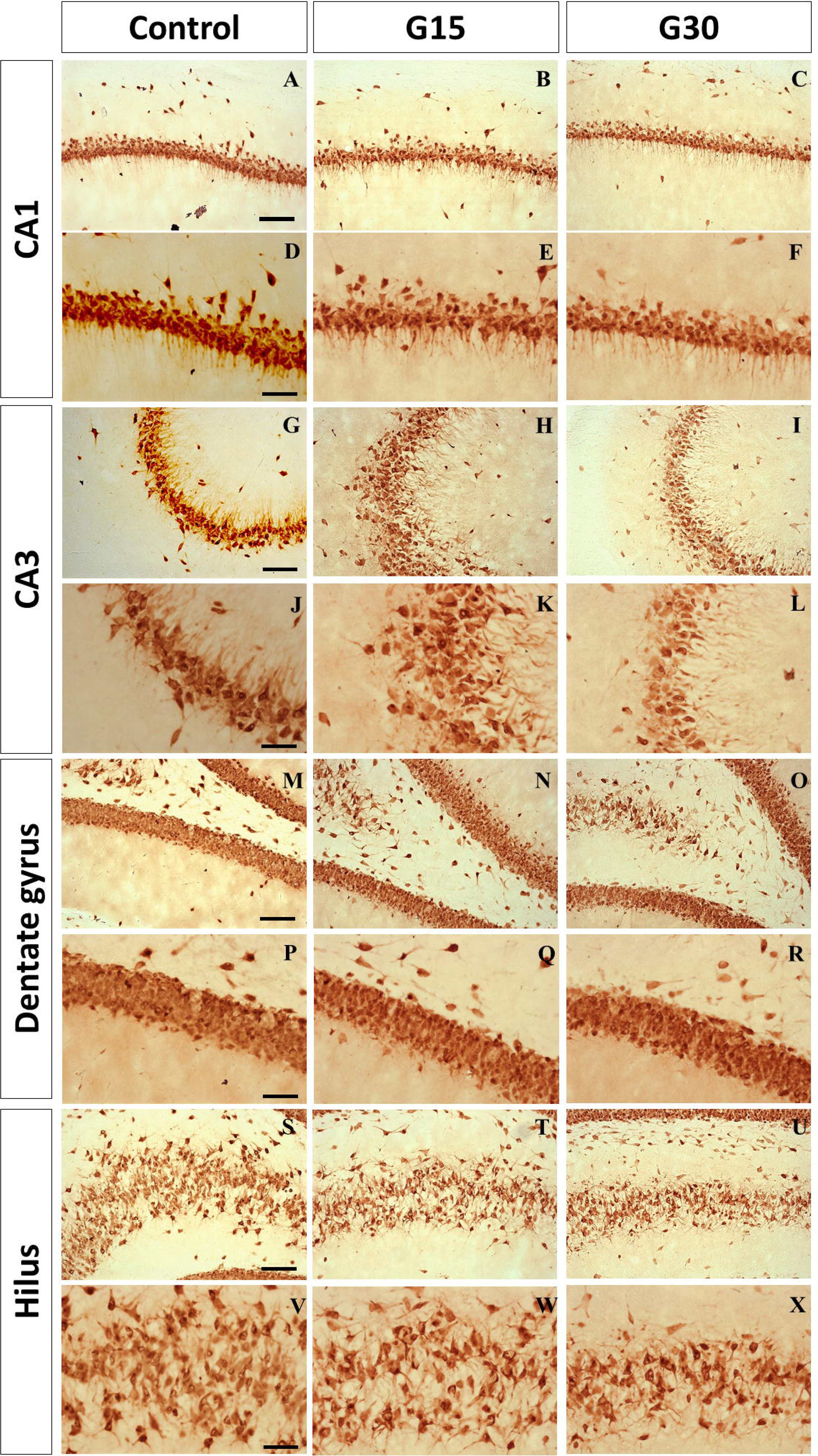
Cytoarchitectonic changes in hippocampal layers after unilateral hippocampectomy. Photomicrographs of hippocampal sections immunolabeled by NeuN at CA1, CA3, dentate gyrus and hilus for all experimental groups. CA1 is not changed fter surgery (A-F). A thicker granule layer and higher numbers of NeuN+ cells were observed in CA3 at G15 (Fig 4I, L), compared to control (Figure 4G, J). In DG, a granule thicker layer was observed at G30. Scale bars: A-B, G-I, M-O, S-U = 100μm; D-F, J-L, P-R, Y-X = 50μm).

In the dentate gyrus (Figures 4M-R), we have noted a thicker granule cell layer in G30 (Figure 4O, R) compared to G15 (Figure 4N, Q), which parallels the increased number of DCX+ cells previously reported for this hippocampal region. In the hilar region, no major differences on morphology and cytoarchitecture were observed (Figure 4S-X).

### 3.4 Behavioral procedures: Exposure to eight-arm radial maze

All animals learned to collect food present at the end of the four arms of the apparatus and no rat presented problems of adaptation to the maze. The maximum time animals remained in the apparatus was 10 minutes (600 seconds), the maximum allowed in the procedure, and the minimum recorded was 143 seconds.

Table 1 shows the percentage of animals that consume all the food present in the apparatus before 10 minutes during the seven days of experiment for all experimental groups. All animals in the control group completed the task from the second day of exposure to the final day, except for day 4, when one animal did not complete (83%). The percentages were lower for the animals in the experimental groups. For G30, the percentage of animals that finished before time was gradually increased, until 100% of the animals finished the test before the time of Day 4 onwards. For Group G15, except for the first day, 83% to 100% of the animals finished the test before 10 minutes.

**Table 1.**
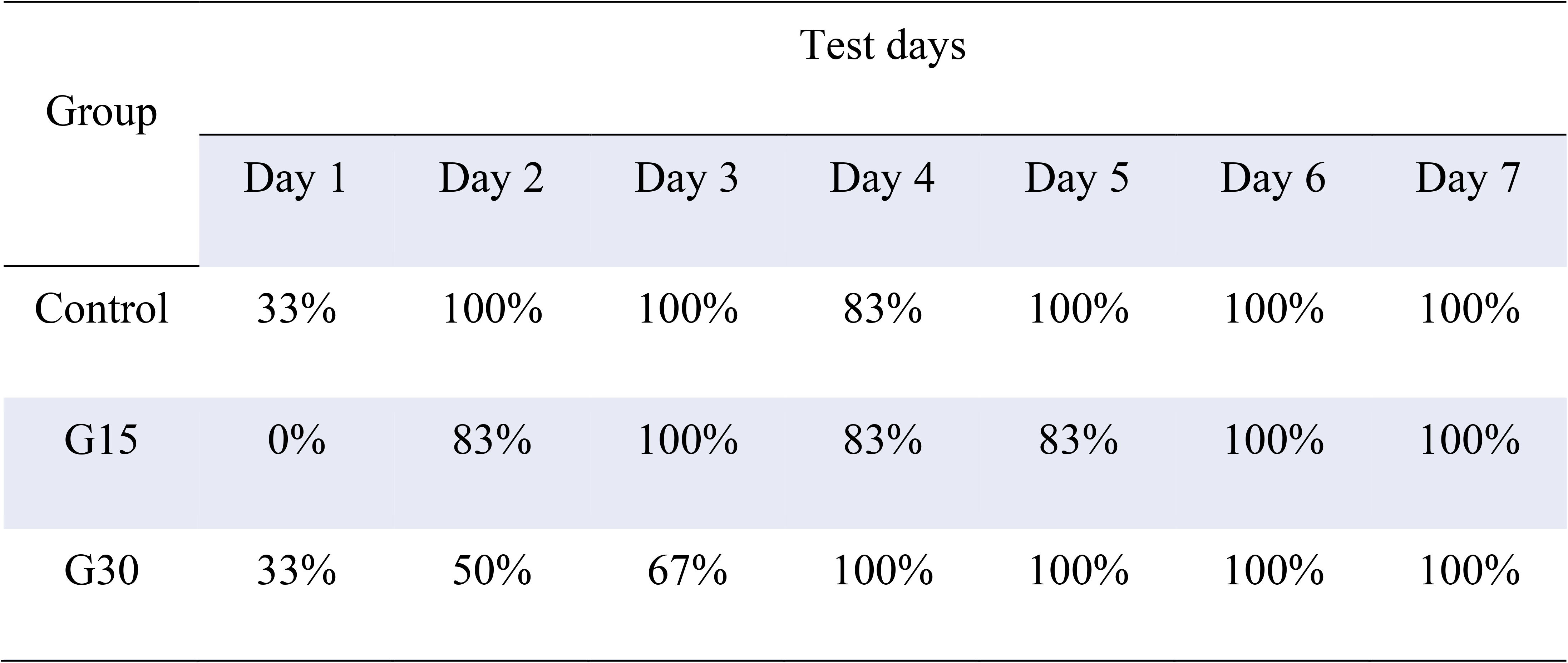
Percentage of animals that completed the eight-arm radial maze task consuming all the food present in the apparatus before 10 minutes in each of the 7-day experiment. The total number of animals exposed per day in each group was 6 (n = 6 per group).

#### 3.4.1 Time of exploration in the eight-arm radial maze

The average time that each group remained in the maze was quantified in all seven days and it was observed a decrease of this time for all groups (Figure 5A) as a function of time, indicating a faster execution of the task, that is, in the consummation of the four pieces of food. Although the shortest time was observed for the control group subjects on all days, comparisons between groups did not present a significant difference (Figure 5A).

**Figure 5.**
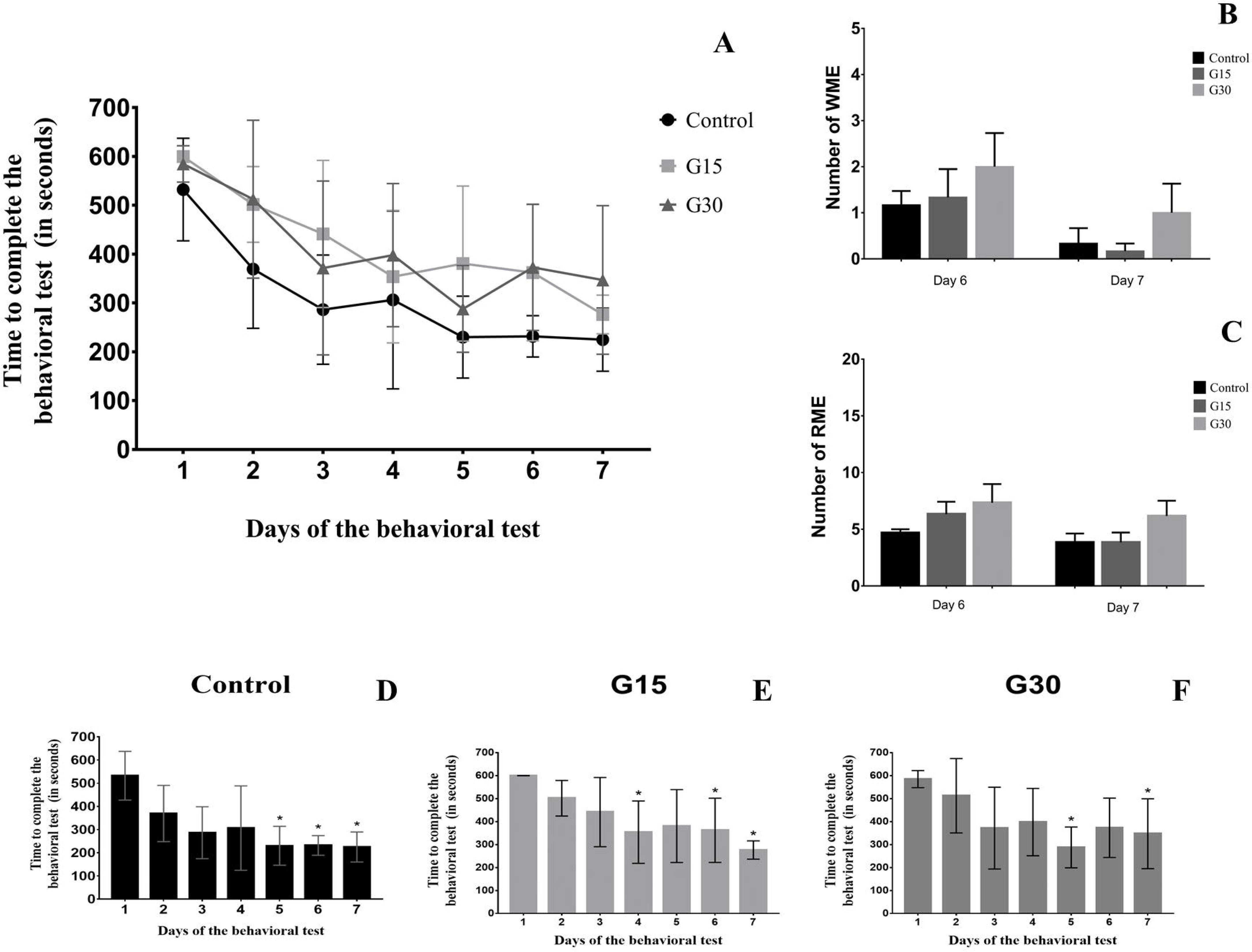
Quantitative data for learning and memory assessment. Average time of exploration of the eight-arm radial maze per group in each experimental day (A). Quantification of working memory errors (B). Quantification of reference memory errors (C). Times for test performance in each experimental day for all groups (D-F). No statistical differences were observed after comparisons (p>0.05, Kruskal Wallis with dunn posthoc-test).

In order to evaluate whether the decrease in intragroup time was significant, we compared the exposure time to the maze in the first day with time in the other days. In the control group, there was a significant difference between the time on day 1 and time on days 5, 6 and 7 (Figure 5D); in G15 group, there was a difference between the time on day 1 and days 4, 6 and 7 (Figure 5E); and in G30 group, there was a difference between day 1 and days 5 and 7 (Figure 5F). Therefore, in general a significant difference was observed in the reduction of time to perform the task between day 1 and day 7 in all groups, control and both surgical.

#### 3.4.2 Hippocampectomy does affect animal performance on working and reference memory behavioral tests

As to the memory measures, the analysis of the working memory errors (WME) and the reference memory errors (RME) of the animals in the maze was performed through the comparison between the control group and the surgical groups on the sixth and seventh day of the experiment. The comparison of the EMT and RME between the groups did not show significant differences on both days analyzed (Figure 5B-C). These results reinforce the idea that learning in the eight-arm radial maze was not permanently affected by surgery.

## 4. Discussion

Unilateral hippocampectomy surgery performed on human patients with refractory TLE promotes the unilateral removal of the hippocampus affected by the pathology as well as the adjacent cortical area (Boling, 2018; Jobst & Cascino, 2015). This surgery is an alternative treatment for those patients who do not get control of seizures with drug treatment and is reported as effective in the control of recurrent epileptic seizures (Liao et al., 2018).

In this study, we developed an experimental model of unilataeral hippocampectomy in adult rats to simulate the human surgery used to treat refractory TLE. The results clearly show that there was a complete removal of the left hippocampus with preservation of the right one. Following surgery, there was an increase in the number of neuroblasts in the contralateral hippocampus at 15 and 30 days post-surgery with morphological alterations in the granular layer at G30, which suggests incorporation of adult granular neurons derived from adult-born neuroblasts. In addition, behavioral tests did not reveal any memory and learning deficits, which parallels the human findings.

### 4.1 The experimental hippocampectomy model is an important achievement for studying compensatory neurogenic events in the hippocampus

The implementation of an experimental model of unilateral hippocampectomy allowed us to study possible neuroplastic events in the contralateral intact hippocampus. This is of clinical significance as simulates the surgery used to treat refractory TLE in humans (Boling, 2018; Jobst & Cascino, 2015). As humans do not present any considerable cognitive impairment after surgery, compensatory neuroplastic events must occur in the contralateral hippocampus. This is likely related to compensatory adult neurogenesis, as hippocampal dentate gyrus is a neurogenic regions in adult brain in both experimental animals (Kempermann, 2015; Lima and Gomes-Leal, 2019) and humans (Boldrini et al., 2018; Moreno-Jimenez et al., 2019; Lima and Gomes-Leal, 2019).

Despite the fact that we have implemented the experimental model of hippocampectomy in non-epileptic animals, this experimental model can be used in animals with chronic epilepsy in future studies. This would simulate better what occurs in human patients with refractory TLE.

### 4.2 Increased adult neurogenesis may occur in the contralateral hippocampus as a compensatory mechanism to avoid cognitive impairment after

Experimental hippocampectomy rendered increased neurogenesis in the contralateral hippocampus up to one month after surgery. This is likely a compensatory neuroplastic event to avoid loss of hippocampal function. This is supported by the fact that animals do not present any important cognitive impairment despite complete unilateral hippocampectomy. The same results are present in humans after unilateral hippocampal removal to treat refractory TLE (Boling, 2018; Jobst & Cascino, 2015).

The mechanisms that triggers compensatory events in the remaining hippocampus are unknown. It has been established that adult neurogenesis is dramatically changed by pathological alterations, including stroke, trauma, epilepsy and other neural disorders (Jin et al., 2006; Lazarov & Marr, 2010; Zhong & Tang 2016). Epilepsy and ischemic stroke induce increase in hippocampal neurogenesis (Koh & Park, 2017; Zhong, & Tang, 2016), while Alzheimer’s induces neurogenesis decrease (Moreno-Jiménez et al., 2019).

We have reported that numbers of neuroblasts were higher in G15 compared to G30 in the rat dentate gyrus. It is possible that these differences are the result of neuroblast maturation at 30 days, which normally induces reduction of DCX expression. The was no difference on the numbers of neuroblast between G15 and G30 in the hippocampal hilus, which suggests a differential pattern of neurogenic events in this region.

In the hilar region, it seems that neuroblast migration continues for a longer time or the level of maturation of adult-born cells is different from the dentate gyrus. The presence of neuroblasts in the hippocampal hilus may represent ectopic migration of adult-born cells to this hippocampal region. This has been reported after status epilepticus (Yang et al., 2008) and stroke (Woitke et al., 2017).

There is a continuous formation of neuroblasts in the adult hippocampus even in elderly humans (Boldrini et al., 2018; Moreno-Jimenez et al., 2019). This precludes an important function for adult neurogenesis on the normal hippocampal function. It has been established that adult-born neurons possess a pivotal role on pattern separation (Sahay et al., 2011), reversal learning and cognitive flexibility, functions related to hippocampal activity (Anacker and Hen, 2017). Loss of these functions may underlie some mood disorders, including anxiety and depression (Anacker and Hen, 2017). The compensatory events here described may play an important role on the avoidance of this pathological events.

### 4.3 Incorporation of adult-born cells into the granule layer may occur in the contralateral hippocampus

We searched for alterations on the normal hippocampal cytoarchitecture after unilateral hippocampal removal. In general, we found no major alterations of density and cellular dispersion between groups. Nevertheless, some changes were present in CA3, as a high density in G15, which was not evident in G30. These differences cannot be explained by incorporation of new neurons, as CA3 is not a neurogenic region. The ectopic migration of granule cells to hippocampal hilus, which connects to CA3 (Meyers et al., 2013), might underlie a transitory rearrange on the CA3 cytoarchitecture, but this deserved further experiments to be confirmed.

In the dentate gyrus, we have noted a thicker granule cell layer in G30, which suggests that adult-born cells may be integrated to into the granule cell layer in the contralateral hippocampus. This is likely a fundamental compensatory event to avoid loss of hippocampal function following unilateral hippocampectomy. Such an event might occur in human patients submitted to hippocampal removal to treat refractory TLE. Future studies, using functional resonance magnetic imaging adapted to visualize human neurogenesis might confirm the present experimental results.

### 4.4 Experimental unilateral hippocampectomy does not induce significative memory deficits, which parallels human surgical patients

The 8ARM allowed the evaluation of inter and intra-group learning by means of measures of the average time spent in the maze and the percentage of animals that completed the task in the course of the days. Seven days of exposure to the maze were used to evaluate learning and memory.

The number of days of exposure to 8ARM varies in the literature, Chan et al. (2004) used twelve days of exposure to the maze, while Kemse et al. (2018) and Rathod et al. (2014, 2015) demonstrated a change in the behavioral patterns of groups evaluated with five days of exposure to 8ARM. In the present study, seven days of exposure appears to have been an adequate choice, as the time was suitable for a minimum survival time (15 days), as well as allowed physical recovery of the animals after the surgical procedure.

Learning and memory are fundamental cognitive functions of organisms and impairments to these functions can significantly affect the quality of life. The reference memory is a type of memory that has a longer duration, and its consolidation is carried out by the hippocampal formation, which can be affected by any damage to this structure; working memory allows the temporary storage of information, being used during the execution of certain tasks and being generally eliminated at the end of the task execution (Chan et al., 2004; Squire & Wixted, 2011).

Anterograde amnesia (inability to consolidate new memories) is described in individuals who have bilateral damage in the temporal lobe area (Squire, 2009; Squire & Wixted, 2011). However, unilateral hippocampectomy surgery is reported to be safe and generally uncorrelated to cognitive impairment related to processing and memory recall to patients (Anyanwu & Motamedi, 2018; Jobst & Cascino, 2015; Mathon et al., 2015; Vojtěch et al., 2012; Wiebe, Blume Girvin, & Eliasziw, 2001). This is in agreement with the experimental results here described, as rats submitted to hippocampectomy did not present any learning and memory deficits.

Considering that hippocampus is an important center of integration and consolidation of memories, the absence of cognitive impairment is likely explained by the fact that the contralateral hippocampus continues to perform its activity, compensating for the absence of the removed hippocampus. Here we propose that compensatory neurogenic events are a fundamental neuroplastic mechanism, which contributes to the absence of learning and memory impairment.

## 5. Conclusion

We conclude that unilateral hippocampectomy induces compensatory neurogenic effect in the contralateral hippocampus. This may underlie the absence of significative cognitive impairment after unilateral hippocampal removal. The results parallels the findings in human patients submitted to unilateral hippocampectomy to treat refractory TLE. These results also establish an experimental model for behavioral evaluation and for analysis of possible compensatory neural processes induced by surgery.

## Acknowledgements

This work was supported by the Brazilian National Council for Scientific and Technological Development (CNPq) and Fundação de Amparo a Pesquisa do Estado do Pará (FAPESPA).

## Data Availability Statement information

The data that support the findings of this study are available from the corresponding author, Silene M.A Lima, upon reasonable request.

## References

Anacker, C., & Hen, R. (2017). Adult hippocampal neurogenesis and cognitive flexibility - linking memory and mood. Nature reviews. Neuroscience, 18(6), 335–346. https://doi.org/10.1038/nrn.2017.45

Anyanwu, C. & Motamedi, G. K. (2018). Diagnosis and Surgical Treatment of Drug-Resistant Epilepsy. Brain sciences, 8. 49. doi:10.3390/brainsci8040049

Boldrini, M., Fulmore, C. A., Tartt, A. N., Simeon, L. R., Pavlova, I., Poposka, V., Rosoklija, G. B., Stankov, A., Arango, V., Dwork, A. J., Hen, R., & Mann, J. J. (2018). Human Hippocampal Neurogenesis Persists throughout Aging. Cell stem cell, 22(4), 589–599.e5. https://doi.org/10.1016/j.stem.2018.03.015

Boling W. W. (2018). Surgical Considerations of Intractable Mesial Temporal Lobe Epilepsy. Brain sciences, 8(2), 35. doi:10.3390/brainsci8020035

Chan, K. F., Jia, Z., Murphy, P. A., Burnham, W. M., Cortez, M. A., & Snead, O. C. (2004) Learning and memory impairment in rats with chronic atypical absence seizures. Experimental Neurology 190. 328–336 doi:10.1016/j.expneurol.2004.08.001

Fisher, R. S., van Emde Boas, W., Blume, W., Elger, C., Genton, P., Lee, P., & Engel, J.Jr. (2005) Epileptic seizures and epilepsy: definitions proposed by the International League Against Epilepsy (ILAE) and the International Bureau for Epilepsy (IBE). Epilepsia. 46. 470–472. doi:10.1111/j.0013-9580.2005.66104.x

Gleeson, J. G., Lin, P. T., Flanagan, L. A. & Walsh, C. A. (1999) Doublecortin is a microtubule-associated protein and is expressed widely by migrating neurons. Neuron 23(2), 257–271.

Jin, K., Wang, X., Xie, L., Mao, X. O., Zhu, W., Wang, Y., Shen, J., Mao, Y., Banwait, S., Greenberg, D. A. (2006). Evidence for stroke-induced neurogenesis in the human brain. Proceedings of the National Academy of Sciences of the United States of America, 103. 198–202. doi:10.1016/j.neuroscience.2015.04.053

Jobst, B. C., & Cascino, G. D. (2015) Resective epilepsy surgery for drug-resistant focal epilepsy: a review. JAMA. 313. 285–293. doi:10.1001/jama.2014.17426

Kempermann, G., Song, H., & Gage, F. H. (2015). Neurogenesis in the Adult Hippocampus. Cold Spring Harbor perspectives in biology, 7(9), a018812. https://doi.org/10.1101/cshperspect.a018812.

Kemse, N., Kale, A., Chavan-Gautam, P., & Joshi, S. (2018) Increased intake of vitamin B12, folate, and omega-3 fatty acids to improve cognitive performance in offspring born to rats with induced hypertension during pregnancy. Food Function. 9. 3872–3883. doi:10.1039/c8fo00467f

Kim, D., Kim, J. S., Jeong, W., Shin, M. S., & Chung, C. K. (2020) Critical area for memory decline after mesial temporal resection in epilepsy patients. Journal of neurosurgery 3, 1–9. doi:10.3171/2019.10.JNS191932

Koh, S. H. & Park, H. H. (2017) Neurogenesis in Stroke Recovery. Translational Stroke Research. 8(1). 3–13. doi:10.1007/s12975-016-460-z.

Lazarov, O., & Marr, R. A. (2010). Neurogenesis and Alzheimer’s disease: At the crossroads. Experimental Neurology, 223, 267–281. doi:10.1016/j.expneurol.2009.08.009

Lima, S., & Gomes-Leal, W. (2019). Neurogenesis in the hippocampus of adult humans: controversy "fixed" at last. Neural regeneration research, 14(11), 1917–1918. https://doi.org/10.4103/1673-5374.259616

Liao, C., Wang, K., Cao, X., Li, Y., Wu, D., Ye, H., Ding, Q., He, H., & Zhong, J. (2018) Detection of Lesions in Mesial Temporal Lobe Epilepsy by Using MR Fingerprinting. Radiology. 288. 804–812. doi:10.1148/radiol.2018172131

Mathon, B., Bédos, U. L., Adam, C., Baulac, M., Dupont, S., Navarro, V., Cornu, P., & Clemenceau, S. (2015) Surgical treatment for mesial temporal lobe epilepsy associated with hippocampal sclerosis. Revue neurologique 171. 315–325. doi:10.1016/j.neurol.2015.01.561

Moreno-Jiménez, E. P., Flor-García, M., Terreros-Roncal, J., Rábano, A., Cafini, F., Pallas-Bazarra, N., Ávila, J., & Llorens-Martín, M. (2019) Adult hippocampal neurogenesis is abundant in neurologically healthy subjects and drops sharply in patients with Alzheimer’s disease. Nature medicine, 25(4), 554–560. doi:10.1038/s41591-019-0375-9

Mullen, R. J., Buck C. R., & Smith A. M. (1992) NeuN, a neuronal specific nuclear protein in vertebrates. Development 116. 201–211.

Paxinos, G. & Watson, C. (1998) The Rat Brain in Stereotaxic Coordinates (Fourth Edition). Press, San Diego, CA.

Rathod, R., Khaire, A., Kemse, N., Kale, A., & Joshi, S. (2014) Maternal omega-3 fatty acid supplementation on vitamin B12 rich diet improves brain omega-3 fatty acids, neurotrophins and cognition in the Wistar rat offspring. Brain and Development. 36. 853–863. doi:10.1016/j.braindev.2013.12.007

Rathod, R. S., Khaire, A. A., Kale, A. A., & Joshi S. R., (2015) Beneficial effects of omega-3 fatty acids and vitamin B12 supplementation on brain docosahexaenoic acid, brain derived neurotrophic factor, and cognitive performance in the second-generation Wistar rats. Biofactors. 41, 261–272. doi:10.1002/biof.1222

Ruan, L., Wang, B., ZhuGe, Q., & Jin, K. (2015). Coupling of neurogenesis and angiogenesis after ischemic stroke. Brain research, 1623, 166–173. doi:10.1016/j.brainres.2015.02.042

Sahay, A., Wilson, D. A., & Hen, R. (2011). Pattern separation: a common function for new neurons in hippocampus and olfactory bulb. Neuron, 70(4), 582–588. doi:10.1016/j.neuron.2011.05.012

Scorza, F. A., Arida, R. M., Naffah-Mazzacoratti Graça, M., Scerni, D. A., Calderazzo, L., & Cavalheiro, E. A. (2009). The pilocarpine model of epilepsy: what have we learned?. Anais da Academia Brasileira de Ciências, 81. 345–365.

Silva, A. F., Aguiar, M. S., Carvalho, O. S., Santana, L. de N., Franco, E. C., Lima, R. R., Siqueira, N. V., Feio, R. A., Faro, L. R. & Gomes-Leal, W. (2013) Hippocampal neuronal loss, decreased GFAP immunoreactivity and cognitive impairment following experimental intoxication of rats with aluminum citrate. Brain Research 1491, 23–33. doi:10.1016/j.brainres.2012.10.063.

Squire, L. R. (2009). The Legacy of Patient H.M. for Neuroscience. Neuron, 61. 6–9.

Squire, L. R., & Wixted, J. T. (2011). The Cognitive Neuroscience of Human Memory Since H.M. Annual Review of Neuroscience. 34. 259–288.

Thored, P., Heldmann, U., Gomes-Leal, W., Gisler, R., Darsalia, V., Taneera, J., Nygren, J. M., Jacobsen, S. E., Ekdahl, C. T., Kokaia, Z. & Lindvall, O. (2009) Long-term accumulation of microglia with proneurogenic phenotype concomitant with persistent neurogenesis in adult subventricular zone after stroke. Glia 57(8), 835–849. doi:10.1002/glia.20810

Vojtěch, Z., Krámská, L., Malíková, H., Seltenreichová, K., Procházka, T., Kalina, M., & Liščák, R. (2012) Cognitive outcome after stereotactic amygdalohippocampectomy. Seizure. 21. 327–333.

Wiebe, S., Blume, W. T., Girvin, J. P., & Eliasziw, M. (2001) A randomized, controlled trial of surgery for temporal-lobe epilepsy. The New England Journal of Medicine. 345. 311–318. doi:10.1056/NEJM200108023450501

Woitke, F., Ceanga, M., Rudolph, M., Niv, F., Witte, O. W., Redecker, C., Kunze, A., & Keiner, S. (2017). Adult hippocampal neurogenesis poststroke: More new granule cells but aberrant morphology and impaired spatial memory. PloS one, 12(9), e0183463. https://doi.org/10.1371/journal.pone.0183463

Yang, F., Wang, J. C., Han, J. L., Zhao, G. & Jiang, W. (2009) Different effects of mild and severe seizures on hippocampal neurogenesis in adult rats. Hippocampus. 18(5), 460–468. doi:10.1002/hipo.20409

Zhong, Q., Ren, B. X., & Tang, F. R. (2016) Neurogenesis in the Hippocampus of Patients with Temporal Lobe Epilepsy. Current Neurology and Neuroscience Reports 16. 20. doi:10.1016/j.expneurol.2009.08.009

